# *HaHB11* transformed maize has improved yield under waterlogging and defoliation in control and field conditions

**DOI:** 10.1101/2021.10.19.465041

**Authors:** Jesica Raineri, Luciano Caraballo, Nicolás Rigalli, Margarita Portapila, María Elena Otegui, Raquel Lía Chan

**Author notes:** **Corresponding author:** Raquel Lía Chan Instituto de Agrobiotecnología del Litoral CONICET-UNL, Centro Científico Tecnológico CONICET Santa Fe, Colectora Ruta Nacional Nº 168 km. 0, Paraje El Pozo. 3000 Santa Fe – Argentina. These authors equally contributed to this work (Luciano Caraballo, Nicolás Rigalli). **Authors’ contributions** Conceived the experiments: JR, MEO, MP, and RLC. JR performed most of the experiments and the illustrations. LC and JR carried out the greenhouse assays. NR and MP performed the spectral reflectance assays and big data analyses and wrote the corresponding items. Conceived and wrote the paper: RLC. JR and MP contributed with the writing. MEO deeply revised and discussed the manuscript. All the authors approved it.

## Abstract

HaHB11 is a sunflower transcription factor previously described as conferring improved yield to maize hybrids and lines. Here we report that transgenic *HaHB11* maize lines exhibited a better performance funder waterlogging, both in greenhouse and field trials carried out during three growth cycles. One of these trials was particularly affected by a strong storm during flowering, causing severe defoliation. Controlled defoliation assays indicated that the transgenic genotypes were able to set more grains than controls. Hybrids were generated by crossing B73 *HaHB11* lines with the contrasting Mo17 lines and tested in the field. These hybrids exhibited the same beneficial traits as the parental lines when compared with their respective controls. Waterlogging tolerance coursed via the root architecture improvement, including more xylem vessels, reduced tissue damage, less superoxide accumulation, and altered carbohydrate metabolism compared to controls. Multivariate analyses corroborated the robustness of the differential traits observed. Furthermore, canopy spectral reflectance data, computing 29 vegetation indices associated with biomass, chlorophyll, and abiotic stress, helped to identify genotypes as well as their growing conditions. Altogether the results reported here indicate that this sunflower gene constitutes a suitable tool to improve maize plants for environments prone to waterlogging and/or wind defoliation.

**One sentence summary:** Phenotyping and big data analyses indicate that the transcription factor HaHB11 confers waterlogging and defoliation tolerance, and increased yield to maize lines and hybrids in all tested conditions.

## Introduction

Maize, rice, and wheat are the crops with the largest production worldwide, providing 60% of the global caloric human intake (FAO 2021, http://www.fao.org/3/u8480e/u8480e07.htm). Maize is used for human nutrition but also for ethanol synthesis and animal feeding. It is a C4 summer crop grown as single-cross (i.e. F1) hybrids presenting high heterosis expression conducive to high grain yield and large biomass production (Duvik, 2005). The improved yield observed in modern hybrids was mainly attributed to enhanced leaf area duration and post-silking crop growth (Rajcan and Tollenaar, 1999), as well as to improved radiation and water use efficiencies (Curin *et al*., 2020). Despite these positive traits, maize is remarkably vulnerable to stress conditions during the critical period of the kernel set (Cerrudo *et al*., 2013) due to the dominated condition within the plant of the grain-bearing organ (the ear) respect to the pollen-producing organ (the tassel). Therefore, most breeding efforts have focused on enhancing abiotic stress tolerance (Chen *et al*., 2016; Tollenaar and Wu, 1999) and reducing apical dominance (Duvik *et al*., 2004; Tollenaar and Wu, 1999).

Although the great efforts devoted to maize breeding, the target production environments of this species are exposed continuously to abiotic stress that penalizes grain yields (Pedersen *et al*., 2017). Among abiotic stress factors and due to global warming, the incidence of floods that expose crops to waterlogging is rising every decade worldwide (Pedersen *et al*., 2017). Flooding events predominate in several areas of the main maize cropping regions. For instance, the May-2018 to April-2019 was the wettest 12-month period in 124 years of records in the United States (NASA, 2019), producing a marked delay in maize sowing date due to soggy soils (USDA, 2019) and a decline in grain yield (FAO, 2019). According to the projection based on multiple climate models, this scenario will not get better; flooding events will increase in most parts of the world during this century (Hirabayashi *et al*., 2021). Global warming also impacts the severity of storms and hail incidence, indicating a significant increase in such phenomena. Strong winds and hail produce different degrees of damage to maize crops depending upon the defoliation intensity and the opportunity of the event (Battaglia *et al*., 2019).

The adaptation of rice to flooding has been deeply studied, being this species resilient to anaerobic soil conditions. The investigation about this harmful stress was divided into that provoked by waterlogging (root system inundation) and the generated by submergence of the aerial system (Bailey-Serres *et al*., 2012a; Voesenek *et al*., 2015). Waterlogging causes quick soil O_2_ depletion because rhizosphere microbes rapidly consume it, provoking changes in the fixation of nitrogen and other nutrients. Another effect of waterlogging is a decrease in soil pH, that increases toxic metals and phosphorous solubility (Setter *et al*., 2009; Bailey-Serres and Voesenek, 2008). In these conditions, plants become unable to cope with evaporative demand, reducing gas exchange and growth (Bramley *et al*., 2007). Gas diffusion is reduced 10^4^-fold, limiting not only oxygen for aerobic respiration but also the CO_2_ needed for photosynthesis (Abiko *et al*., 2012).

When plants sense these environmental changes, they trigger molecular signaling pathways to cope with the stress, including specific modulation of gene expression and hormone homeostasis (Voesenek and Bailey-Serres, 2015). Among the hormones involved in plant response to flooding, ethylene plays a key role (Bailey-Serres *et al*., 2012b; Bailey-Serres and Voesenek, 2010; Loreti *et al*., 2016; Sasidharan *et al*., 2018; Voesenek and Sasidharan, 2013). Ethylene accumulates in the cells, eliciting the formation of reactive oxygen species, which play a dual role as signaling molecules and causing oxidative stress damage (Sasidharan *et al*., 2018; Yamauchi *et al*., 2018).

Adaptation to waterlogging stress also involves the fine-tuning of several genes, mostly associated with carbohydrate transport, anaerobic metabolism, cell wall remodeling, and detoxification. Among these genes, there are those encoding the enzymes invertase (*INV*); glucose-6-phosphate isomerase (*G6PI*); glyceraldehyde-3-phosphate-dehydrogenase (*GAPDH*); phosphoglycerate kinase (*PGK*), and alcohol dehydrogenase (*ADH*) (Arora *et al*., 2017; Du *et al*., 2017; Zou *et al*., 2010). The root system is the most affected by waterlogging, and the adaptive changes include the generation of new adventitious roots and aerenchyma constituting a barrier to radial oxygen loss (Abiko *et al*., 2012, Loreti *et al*., 2016; Yamauchi *et al*., 2018).

Regarding defoliation stress, it is caused by varied factors such as storms, hail, leaf diseases, and herbivore attacks, causing all them a total or partial reduction in the leaf area of plants, leading frequently to reduce light interception, and consequently less biomass production through photosynthesis. In early defoliation events, the grain yield penalization is usually low provided the apical meristem is not injured causing plant death and stand reduction (Battaglia *et al*., 2019). By contrast, partial defoliation during the critical period for kernel set may decrease seed yield dramatically (Battaglia *et al*., 2019), depending upon the reduction caused in plant growth rate (Andrade *et al*., 1999). The extent of yield penalization due to defoliation during the active grain-filling period will depend upon the reduction caused to the source-sink ratio during this stage (Borrás *et al*., 2004).

Environmental factors are perceived by plants that display signal transduction pathways resulting in the degradation of superfluous biomolecules and the synthesis of others needed to deal with stress. In the first steps of such molecular responses, transcription factors (TFs) play a crucial role as master switches able to activate or repress entire metabolic pathways. In plants, there are numerous TFs (more than 1500 in the model Arabidopsis) classified in families, mainly according to the conserved DNA binding domain. Among these families, the homeodomain-leucine zipper (HD-Zip) is unique to this kingdom and was associated with abiotic stress responses (Perotti *et al*., 2017). Members of this family present high conservation of the HD-Zip domain, and the sequencing of whole genomes of different species revealed other uncharacterized functional motifs located in the N- and C-termini of these proteins (Arce *et al*., 2011). Notably, in sunflower and other Asteraceae species, there are HD-Zip I proteins exhibiting distinctive carboxy-termini. Among these divergent members, there is HaHB4, which confers tolerance to drought in wheat and soybean plants (González *et al*., 2020), and HaHB11, which enhanced yield in B73 lines and HiII hybrids (Raineri *et al*., 2019). HaHB11 also conferred flooding tolerance to Arabidopsis plants, both to waterlogging and submergence (Cabello *et al*., 2016).

*HaHB11* maize plants were assessed in greenhouse and field trials during three growing seasons. Phenotyping was carried out by measuring different traits conducive to characterize plant and crop growth, such as stem width and height, leaf area, total biomass, ASI (anthesis-silking interval), light interception, and grain yield (Raineri *et al*., 2019).

In this work, we describe greenhouse and field trials revealing that maize plants expressing the sunflower TF HaHB11 exhibit enhanced tolerance to waterlogging. Moreover, during one of these trials, a strong storm that provoked severe defoliation revealed that transgenic plants were able to withstand the negative effects of defoliation better than the non-transgenic ones. This response was corroborated in subsequent controlled assays. We obtained new hybrids, using transgenic and non-transgenic B73 lines crossed to the contrasting MO17 parental. Transgenic hybrids had increased yield compared to the controls. Finally, non-destructive spectral analysis (remote-sensing) along the cycle of field-grown crops allowed distinguishing controls from transgenic genotypes.

## Results

### *HaHB11* transgenic maize plants exhibit increased tolerance to waterlogging compared with controls in greenhouse and field assays

Although the enormous differences between the sunflower (the species from which *HaHB11* was isolated) and the model plant Arabidopsis, the evolutionary distance between Arabidopsis and maize is even greater. Hence, we wondered if the waterlogging tolerance, conferred by HaHB11 to Arabidopsis, was conserved in maize.

Firstly, we carried out waterlogging assays with plants grown on pots filled with sand in the greenhouse. Several characteristics were assessed in two independent transgenic lines and null segregants, used as controls (B73). During the treatment period, transgenic plants developed longer roots with increased biomass and achieved a larger leaf area than controls (Figures 1A, 1B, 1D, 1E). Moreover, compared with the null segregants, the leaves of *HaHB11* plants showed higher stomatal conductance (Figure 1C). Overall, these results suggest an increased waterlogging tolerance of *HaHB11* plants.

**Figure 1.**
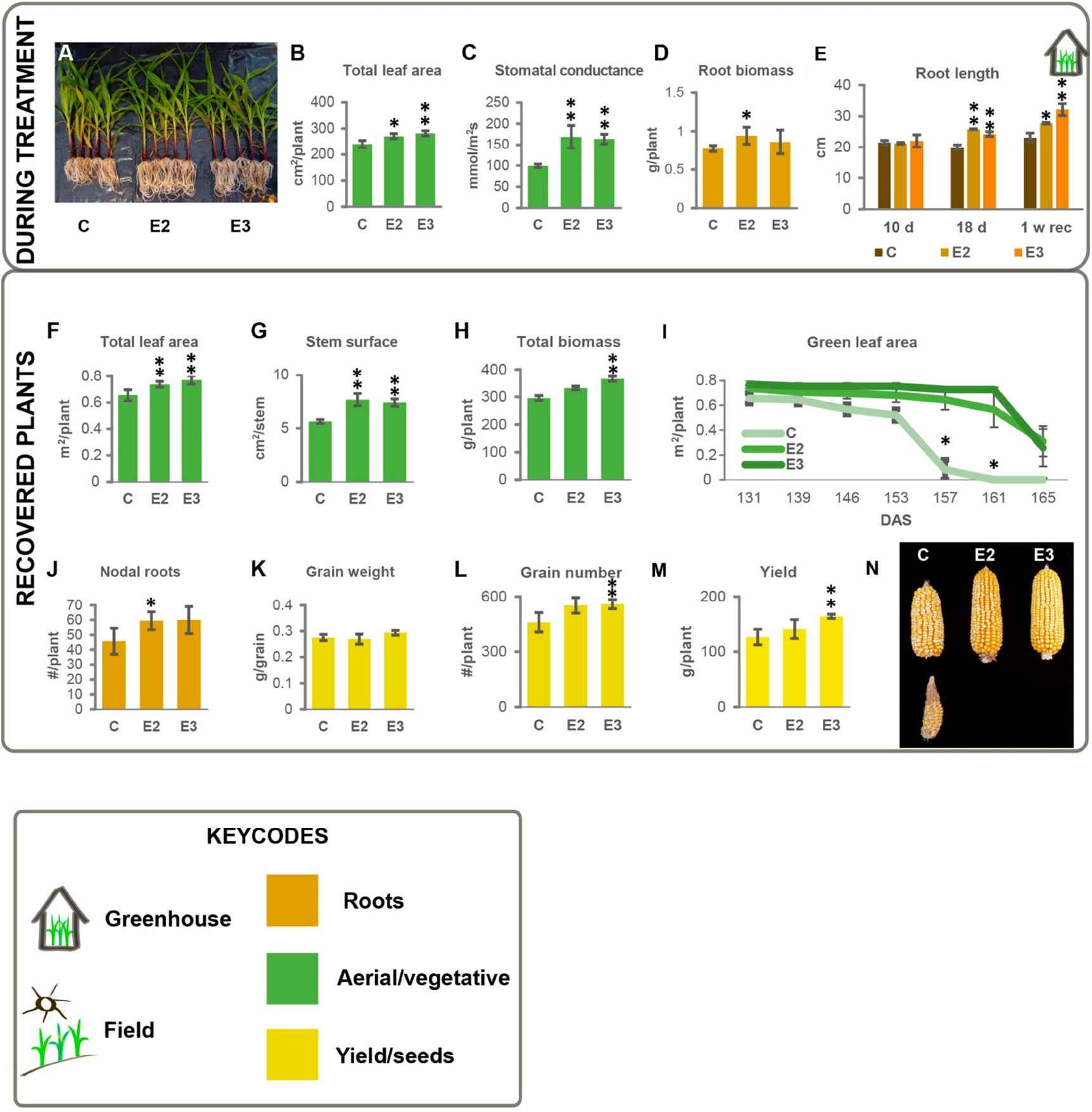
Transgenic plants expressing *HaHB11* exhibit enhanced waterlogging tolerance compared with their B73 controls in the greenhouse. Illustrative picture of maize plants subjected to waterlogging during 18 days (A). Total leaf area (B) and root biomass (D) after 18 days of waterlogging. Stomatal conductance after 7 days of treatment (C). Root length after 10 and 18 days of waterlogging treatment plus 1 week of recovery (E). Total leaf area at silking (F), stem surface (G), total biomass (H), green leaf area (I), number of nodal roots (J), grain weight (K), grain number (L) and yield (M). Illustrative pictures of cobs at harvest of control (C), and transgenic events (E2 and E3) in B73 background (N). Data represent means ± SEM of at least 4 biological replicates. Asterisks indicate significant differences respect to the control genotype (* for P<0.05 and ** for P<0.01). Bottom: codes used in all the illustrations.

In the field, waterlogging usually persists for several days but not along all the life cycle. Hence, after the stress treatment, plants were placed into larger pots and were allowed to recover and grow in normal conditions. At the end of the life cycle, *HaHB11* transgenic plants developed larger leaf and stem areas and an extended period of leaf greenness and concurrent delayed senescence (Figures 1F-1I). These trends were accompanied by increased biomass (Figure 1H). Moreover, such plants had more nodal roots (Figure 1J) and enhanced grain number (Figure 1L), whereas individual grain weight (Figure 1K) did not differ between genotypes. Described traits explained the increased grain yield (Figure 1M) and healthy aspect of the produced kernels (Figure 1N). Notably, most differential traits between controls and transgenics, observed in normal conditions trials (Raineri *et al*., 2019) were maintained after this stress treatment.

Even though greenhouse assays gave us a preliminary idea about the performance of *HaHB11* transgenic plants after a waterlogging episode, field-grown maize is exposed to a combination of environmental conditions (irradiance level, wind, evaporative demand, etc.) that may modify results obtained in the greenhouse. However, the generation of waterlogging conditions in the field is rather difficult. Hence, we designed a mixed test to evaluate the performance of waterlogged maize (see Methods). Waterlogging was applied for 14 days to V4 plants grown in a large pot in the field (Figure 2A). After that, plants were transferred to soil and grown in standard conditions until harvest (Figure 2A). To further understand the distinctive root phenotype observed in the greenhouse (Figures 1A, 1D-E), we performed and analyzed transversal cuts. This study indicated that the transgenic genotype developed more xylem vessels than controls (Figures 2B, 2C). Similar to the greenhouse scenario, *HaHB11* plants exhibited delayed senescence and increased chlorophyll content than controls (Figures 2D, 2E, 2F), and developed more nodal roots, wider stems, and total aerial biomass (Figures 2G, 2H, 2I), indicating a better recovery from waterlogging than their control counterparts. Regarding grain yield determination, transgenic plants partitioned more biomass into grains (Figure 2J), showing increased yield, explained by an improved grain setting (Figures 2L, 2K).

**Figure 2.**
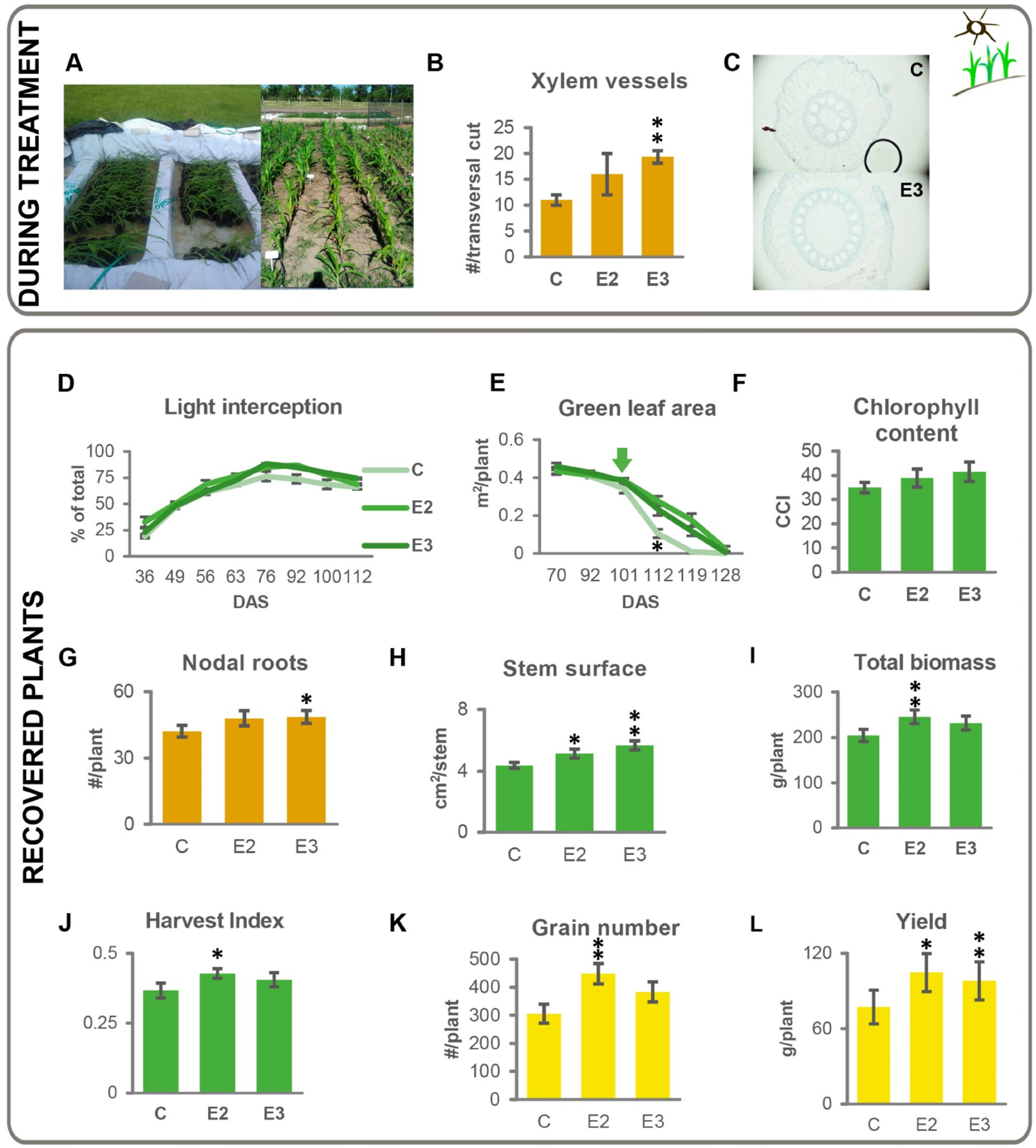
Field-grown transgenic plants expressing *HaHB11* tolerate better waterlogging than their B73 controls. Illustrative picture of waterlogging treatment in the field (left, A) and plants placed on soil after treatment (right, A). Xylem vessels per transversal root cuts (B) and image of the stained cross sections of control and E3 transgenic plants (C), after 2 weeks of waterlogging. Light interception and green leaf area during the life cycle (D, E). Ear leaf green leaf index, 100 days after sowing measured as chlorophyll/carotenoid index (CCI, F). Nodal roots number (G), stem surface (H), total biomass (I), harvest index (J), grain number and yield (K, L) of plants at harvest. Data represent means ± SEM of at least 3 biological replicates. Asterisks indicate significant differences respect to the control genotype (* for P<0.05 and ** for P<0.01).

### *HaHB11* transgenic maize plants withstood the hardship of an unexpected severe windstorm and exhibited improved performance than controls under defoliation

During the third waterlogging assay in the field, a severe windstorm with gusts of 107 km h^-1^ hit the crop eleven days after silking, when still in the critical period for grain setting. Plants that were not killed and remained standing were completely defoliated with leaves preserving only their midribs (Supplementary Figure S1). Surprisingly, transgenic *HaHB11* plants accumulated more biomass and doubled grain yield of controls at maturity (Supplementary Figure S1). To confirm this serendipitous finding, we performed a greenhouse assay. We manually defoliated the plants 11 days after silking with similar results to those of the field (Supplementary Figure S2). These results strongly suggested that transgenic plants can withstand defoliation during seed filling better than the controls, and therefore reduced the associated penalization to grain yield.

### Is HaHB11 able to improve the already enhanced growth promoted by heterosis in F1 maize hybrids?

Original transgenic maize plants were obtained in the HiII hybrid (a cross between the A188 and B73 inbreds) of poor performance compared with commercial hybrids. Hence, the progeny of HiII was backcrossed to B73 to recover the phenotype of this inbred line and reduce phenotypic segregation (Raineri *et al*., 2019). The beneficial effect of *HaHB11* on several agronomic traits was detected, albeit at different extents, dependent on the heterozygosity levels.

Heterosis in maize usually increases yields around 72-254% under no-stress conditions (Duvik, 2005; Munaro *et al*., 2011). Thus, we wondered if *HaHB11* would be able to maintain the previously described beneficial traits when expressed in an improved hybrid background, or the benefits conferred by the transgene may be masked due to the enhanced heterosis conferred by the cross of inbreds representative of contrasting heterotic groups. We performed crosses between B73 (transgenic and control plants) and the Mo17 public lines. The former belongs to the Reid Yellow Dent Group and the latter to the Lancaster Sure Crop Group, and crosses between them have been widely studied (Troyer, 1999). In greenhouse assays, carried out in normal growth conditions, transgenic F1 hybrids, from the B73 × Mo17 cross, accumulated more biomass, and exhibited delayed senescence (Figures 3A, 3B, 3C). Moreover, similar to the results obtained with lines, transgenic hybrids achieved significant higher grain number and yield than controls (Figures 3D, 3E). These results strongly suggested that *HaHB11* expression could still improve hybrid plants, at least in standard conditions.

**Figure 3.**
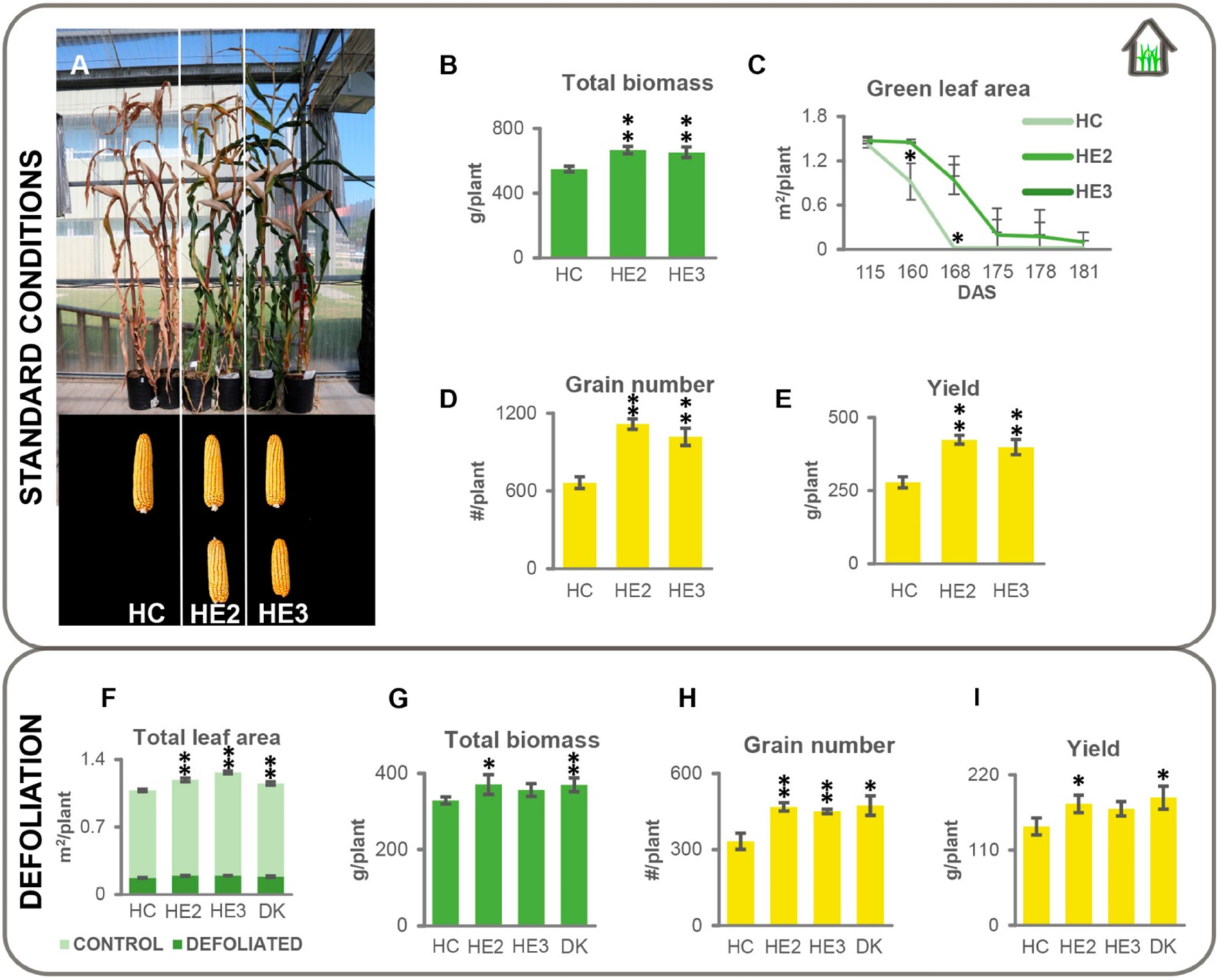
*HaHB11* transgenic hybrids tested in the greenhouse, exhibited delayed leaf senescence and greater yield than controls in standard growth conditions and after defoliation. Upper panel: Illustrative photograph of B73 x Mo17 hybrids plants and their cobs (A). Total biomass (B). Senescence during grain filling (C). Grain number (D) and yield (E) at the end of life cycle. Plants were grown in normal conditions. Lower panel: Total leaf area of plants before (F, light green), and after defoliation (F, dark green). Total biomass, grain number and yield of defoliated plants at harvest (G, H, I). Data represent means ± SEM of at least 4 biological replicates. Asterisks indicate significant differences respect to the control genotype (* for P<0.05 and ** for P<0.01).

### *HaHB11* transgenic hybrids exhibited enhanced tolerance to waterlogging and defoliation in greenhouse and field assays

Aiming at knowing *HaHB11* hybrids performance under abiotic stress conditions, we carried out defoliation and waterlogging assays firstly in the greenhouse. Defoliation was performed manually 11 days after silking on transgenic and control hybrids. The leaf area removed from *HaHB11* plants was slightly larger than from controls because individual leaf area was larger among plants of the former (Figure 3F). Despite defoliation, at the end of the cycle, transgenics had increased biomass, seed yield, and grain number than control plants (Figure 3G, 3H, 3I). These results were similar to those obtained with the parental line B73, both in the field and the greenhouse. Notably, the F1 hybrid DK72-10® included as a reference, for comparison with a commercial product currently used by farmers, showed similar results as *HaHB11* ones.

Regarding waterlogging, root development was assessed by measuring different traits related to flooding tolerance. Similar to *HaHB11* lines, transgenic hybrids exhibited increased root volume and biomass, as well as a higher number of xylem vessels/pith area than controls (Figures 4A, 4B, 4C, 4E). To understand whether these differential traits lead to differences in radial oxygen loss, we treated the roots with methylene-blue. Figure 4D confirmed that control roots lost more oxygen (blue-stained roots) than *HaHB11* roots, indicating that this mechanism could be contributing to the hypoxia tolerance showed by the transgenic plants. This result may explain the enhanced chlorophyll content (Figure 4F) and the number of vascular bundles (Figures 4G, 4H) of transgenic plants with respect to controls detected on 14 days after waterlogging when plants were already growing in standard conditions. At flowering, leaves were larger in the transgenics than in the control plants (Figure 4I), and both independent events exhibited delayed senescence (Figure 4L). Moreover, the transgenics showed a higher number of nodal roots, compared with controls on the hybrid background (Figure 4J, 4K). All these characteristics explained, at least in part, the increased total biomass and grain yield of *HaHB11* plants (Figure 4M, 4N).

**Figure 4.**
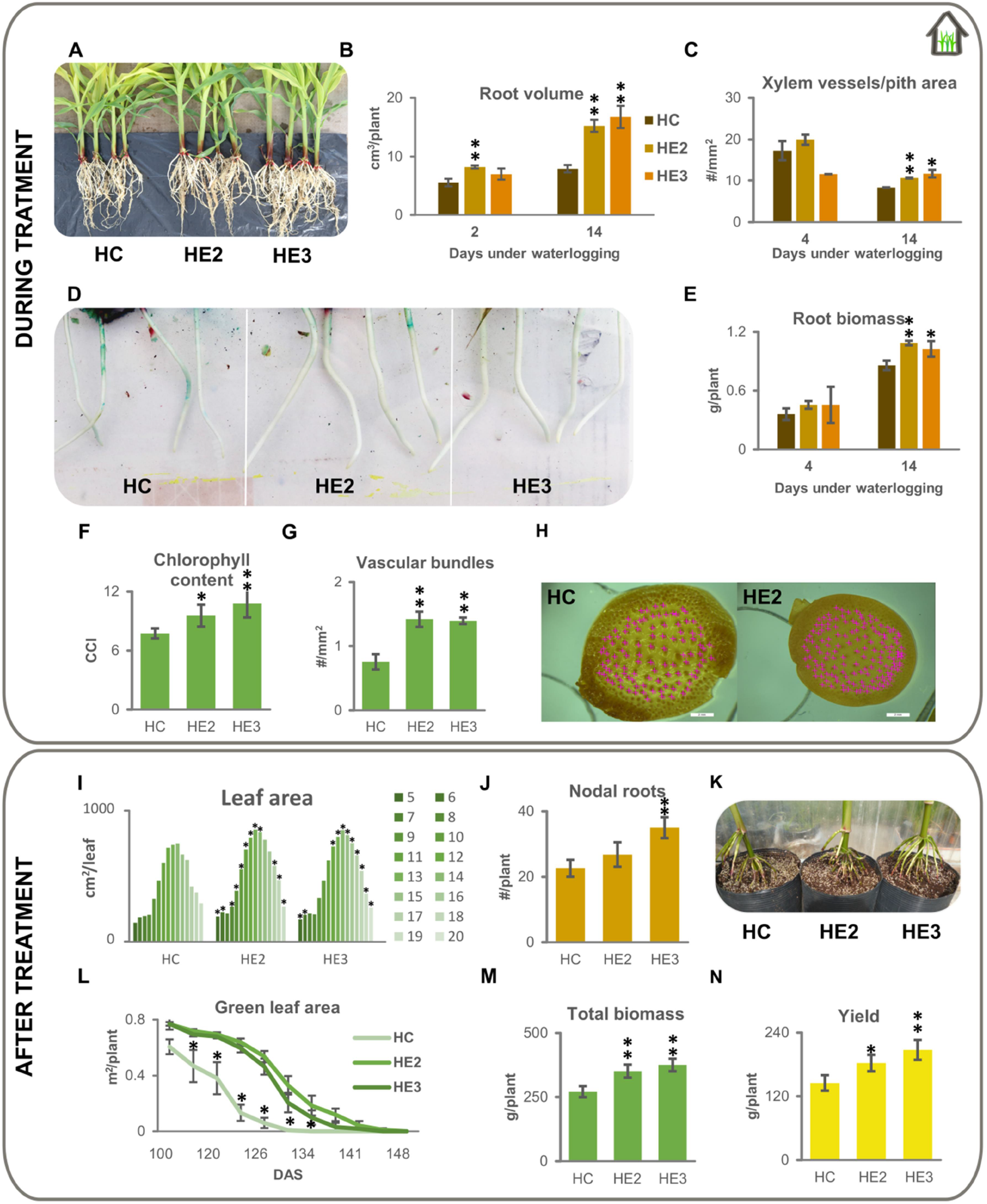
Transgenic hybrids expressing *HaHB11* showed improved performance during waterlogging stress and after recovery than controls in the greenhouse. Illustrative image of hybrid controls (HC) and transgenic events (HE2 and HE3) after two weeks of waterlogging treatment (A). Root volume (B), xylem vessels per pith area (C), and root biomass (E). Illustrative photograph of roots stained during one hour with methylene blue, taken 2 weeks after initiating waterlogging treatment (D). Chlorophyll content of the 5^th^ leaf after 14 days of waterlogging treatment (F). Illustrative picture of vascular bundles in transversal sections of stems. Vascular bundles are marked with a pink cross (G, H). Phenotype of maize plants subjected to 14 days of waterlogging, and then grown in normal conditions until harvest. Individual leaf area from leaf 5 to 20 (I). Total nodal roots and representative photograph of the plants (J, K). Green leaf area of plants at the end of grain filling (L). Total biomass and yield at harvest (M, N). Data represent means ± SEM of at least 4 biological replicates. Asterisks indicate significant differences respect to the control genotype (* for P<0.05 and ** for P<0.01).

In the field, the results were similar to those observed in the greenhouse when roots were evaluated after 12 days of waterlogging treatment. In transversal cuts of adventitious roots, transgenic plants developed a higher number of xylem vessels/pith area (Figure 5A, 5D). Tissue damage was detected on the pith of control roots, whereas the medulla was intact in the transgenic ones (Figure 5A). Such damage could be generating an impaired function in control roots. Moreover, transgenic roots accumulated more lignin than the control ones (Figure 5B), suggesting that controls lost more radial oxygen than transgenics. Stating that transgenic plants deal better with the oxidative stress triggered by waterlogging, NBT staining was carried out seven days after the treatment, resulting in reduced superoxide accumulation in *HaHB11* plants (Figure 5C).

**Figure 5.**
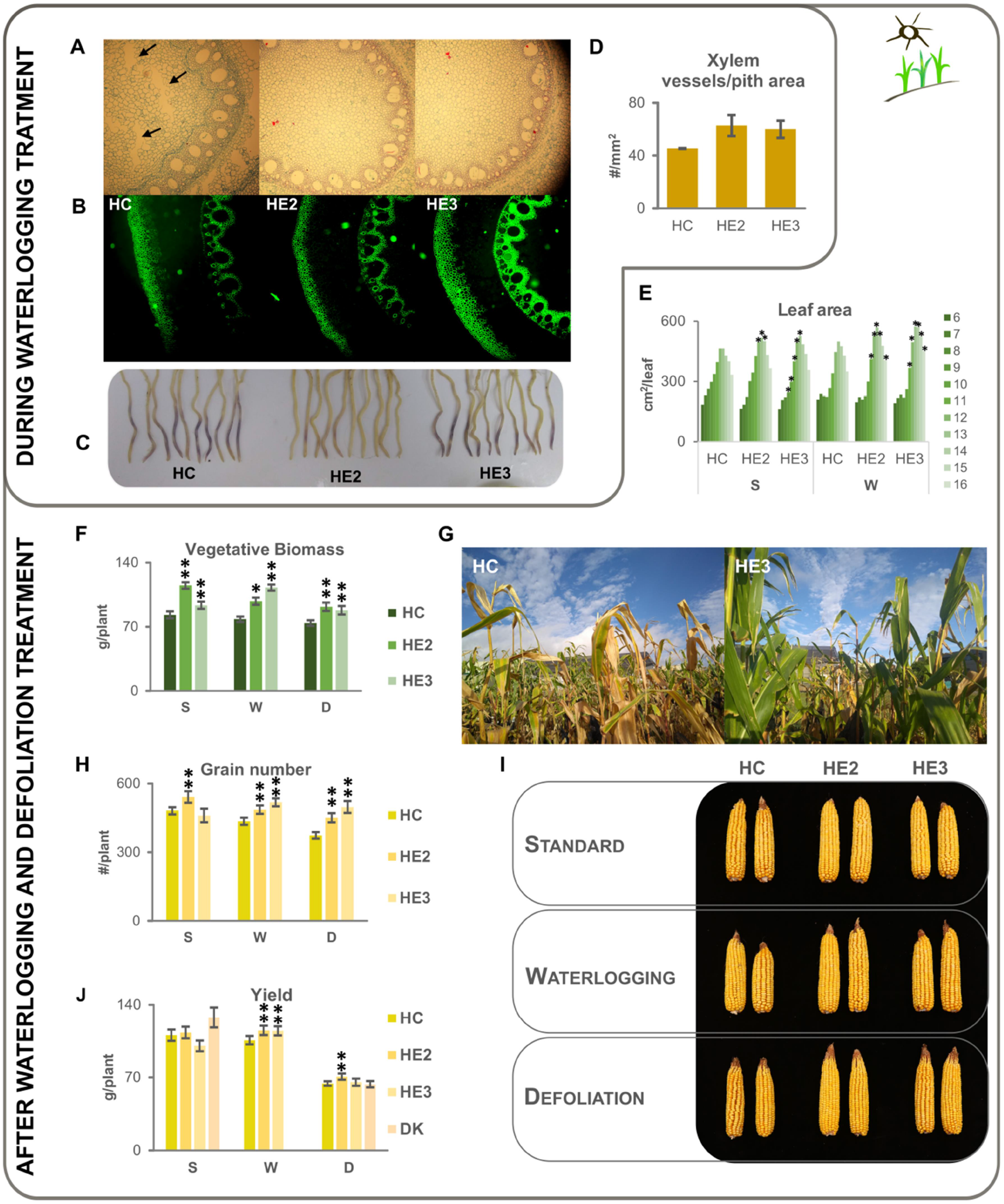
Field-grown transgenic hybrids carrying *HaHB11* exhibited a better performance than their controls grown in standard conditions and after waterlogging or defoliation treatments. Transversal cuts of adventitious roots, 12 days after initiating waterlogging treatment; samples were taken at 4.5 cm from the root tip (from 6-8 cm length roots), stained with safranin-fast green and captured with white light (A), or epifluorescence (B). Illustrative picture of roots on 7 days after the waterlogging treatment, stained with NBT (C). Number of xylem vessels per pith area in stems of plants waterlogged during 14 days (D). Phenotype of plants grown in normal conditions (S), defoliated 15 days after silking, or (D) subjected to 14 days of waterlogging (W). Individual leaf area (from leaf 6 to 16, E). Vegetative biomass, grain number and yield of plants at harvest (F, H, J). Illustrative photograph of control hybrids c (HC) and two transgenc events (HE2, HE3) on 100 days after sowing (G). Representative picture of ears (HC, HE2, HE3) after different treatments (I). Arrow heads in A point at tissue damage on the pith. Data represents means ± SEM of at least 3 biological replicates. Asterisks indicate significant differences respect to the control genotype (* for P<0.05 and ** for P<0.01).

Once the plants were growing in standard field conditions, the phenotype was assessed until the end of the life cycle. Individual leaf area was larger in *HaHB11* recovered plants compared to controls, and as in other mentioned assays, they had delayed senescence and developed more biomass (Figures 5E, 5F, 5G). Grain number and yield were higher for *HaHB11* plants in control conditions as well as after defoliation and waterlogging (Figure 5I, 5J). The results suggest that yield increase is mainly due to the enhanced grain number (Figure 5H). As expected, the DK72-10 hybrids yielded more than the rest of the evaluated genotypes in standard growth conditions (Figure 5J). However, penalization after defoliation was the highest for this hybrid, and its seed yield was similar to that of transgenic *HaHB11* plants (Figure 5J).

### *HaHB11* modulates the expression of genes involved in carbohydrate metabolism, detoxification, and waterlogging response

To unravel the molecular basis of waterlogging tolerance exhibited by *HaHB11* plants, we selected genes described as differentially regulated in tolerant accessions or after waterlogging treatments.

Transcriptional regulation is a dynamic and fine-tuned process that changes depending on various factors such as stress conditions. We were particularly interested in gene expression kinetics in *HaHB11* plants modulated by waterlogging. Samples were harvested from roots of plants grown in the greenhouse one day after treatment initiation, and from roots and leaves of field-grown plants on 4, 6, and 12 days after treatment initiation.

In roots, expression levels of *GAPDH, ADH, G6PI, BE7, PG, SUT1, AP2, GLK1, HMT, HQX, MS1, PGK*, and *INV2*, mainly involved in carbohydrate metabolism and transport, were assessed. In the greenhouse, *G6PI, GAPDH, INV2, and AP2* were differentially induced in *HaHB11* plants, whereas *ADH, PGK, MS1*, and *HMT* showed the opposite regulation (Figures 6 and S3). The scenario changed in the field. In these conditions, the more remarkable results were the earlier induction of *BE7* in the transgenics (4 days) compared to controls (6 days), and the faster repression of *GADPH* on 4 days after treatment (Figure 6). Among the selected genes, several did not show differential regulation between genotypes, and others were not detectable in roots or leaves harvested at these developmental stages (Supplementary Figure S3).

**Figure 6.**
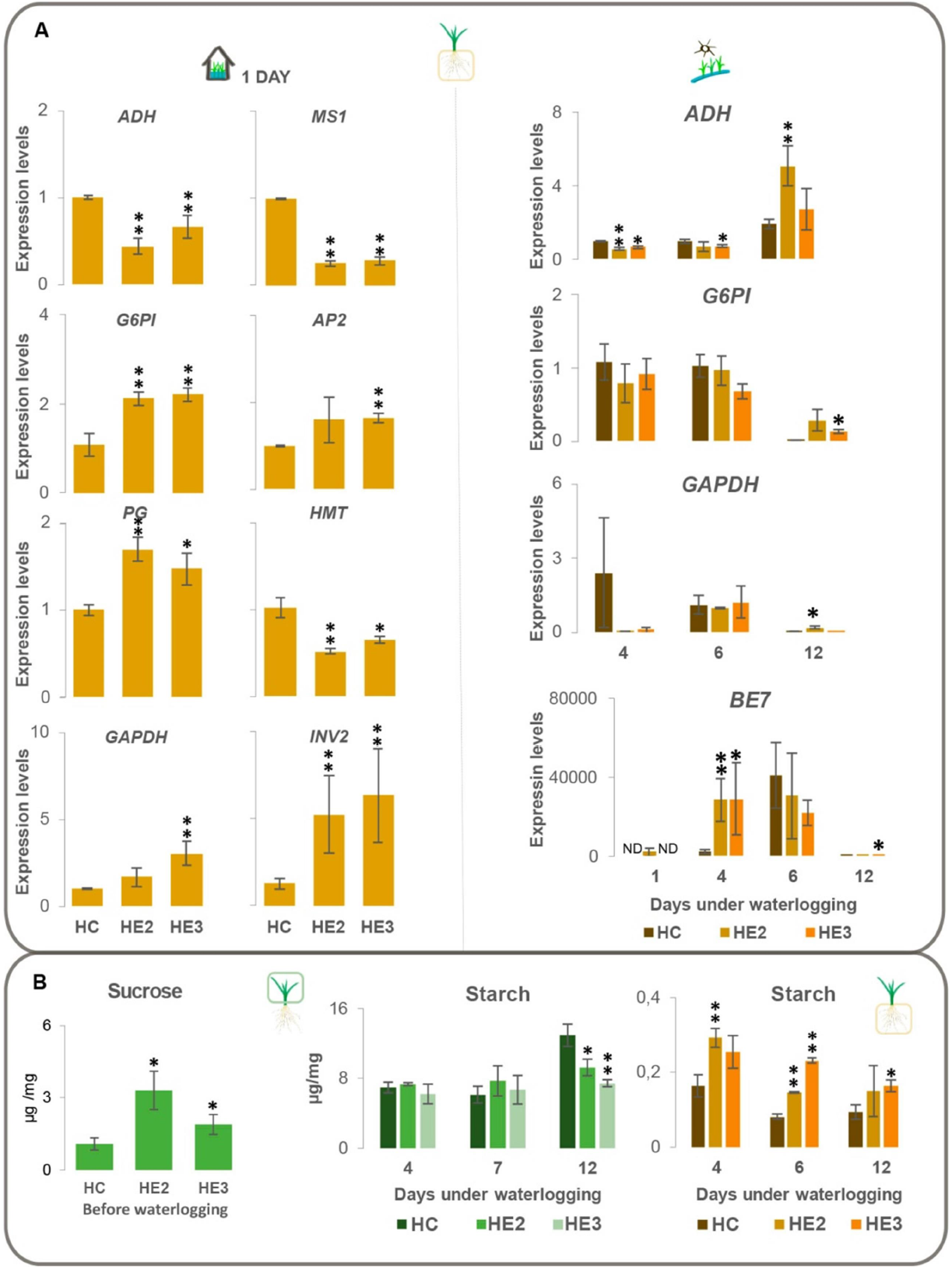
Transgenic HaHB11 hybrids showed differential expression of genes related with waterlogging tolerance and carbohydrate contents. A. Transcript levels of *ADH, G6PI, GAPDH, INV2, PGK, MS1, AP2*, and *HMT* genes involved in waterlogging response in *HaHB11* transgenic roots of the control (HC) and transgenic hybrids (HE2 and HE3), evaluated one day after initiating waterlogging treatment (left panel), or after 4, 6, and 12 days of treatment (right panel). B. Sucrose and starch contents before or after waterlogging treatment in leaves and roots of the control (HC) and transgenic hybrids (HE2 and HE3). All the values were normalized with endogenous *ACTIN* and then with the one obtained in the control, arbitrary assigned a value of one. The ID codes for the tested genes are listed in Supplementary Table S1. Each point is the average of four plants and error bars represent standard error of the mean (SEM) x 2. Asterisks indicate significant differences respect to the control genotype (* for P<0.05 and ** for P<0.01).

In leaves, the evaluated genes were *GAPDH, ADH, GPI, SWEET13*, and *ACS3*. Such genes are involved in carbohydrate metabolism and transport. Among them, those showing significant differences between genotypes were *GAPDH, ADH*, and *G6PI. GAPDH* was only induced in *HaHB11* plants after 12 days of treatment (Figure 6A), whereas *G6PI* showed higher expression levels in the transgenic leaves than in the control ones, both after 4 or 12 days of treatment (Figure 6A). Other evaluated genes did not show differential regulation between genotypes or treatments (Supplementary Figure S3).

To assess whether the observed transcriptional changes affected carbohydrate contents, sucrose and starch concentrations were evaluated in roots and leaves of waterlogged plants. In leaves, the hybrids HE2 and HE3 had more sucrose than controls. After 4 and 7 days of waterlogging, starch content was similar in all the genotypes; however, 12 days after initiating the treatment, controls accumulated more starch in the leaves than transgenic hybrids, suggesting that *HaHB11* plants were more efficient to deliver carbohydrates to other active growing sinks. In agreement, roots of *HaHB11* hybrids exhibited more starch than controls after 4, 7, and 12 days of treatment (Figure 6B).

### *In situ* canopy spectral reflectance helped discriminate maize genotypes nondestructively across growing conditions

Traditional ground-based crop phenotyping of secondary, physiological traits aimed at breeding is usually limited by the number of plants that can be evaluated and is being replaced by nondestructive methods such as spectral images (Reynolds *et al*., 2021), which produce a large number of data and consequently the need of adequate computing tools for their analysis. Evaluation of canopy spectral reflectance in crops is done predominantly by studying a collection of vegetation indices (VIs), comparing their performance to select a single one or a few of them that better represent a trait of interest (Reynolds *et al*., 2021). It is less customary to explore the potential of several VIs jointly for capturing differences among genotypes within a single environment (Arias *et al*., 2021), as we did in current research using a set of 29 selected VIs to discriminate maize genotypes grown under different conditions.

Statistical significances for VI values were analyzed for the factor genotype with a one-way analysis of variance (ANOVA) and a posthoc Tukey test. We chose a p-value of 0.05 for statistical significance. Each ANOVA was performed on a set of 162 VI values, comprising 18 measurements per plot on the nine plots per treatment (three repetitions for each of the three genotypes). One ANOVA per date and treatment was performed, giving a total of 261 (three dates, three treatments, and 29 VIs) ANOVAs for VI and their corresponding posthoc tests. When the spectral behavior of the evaluated genotypes was analyzed, we detected that the set of VIs that allowed their discrimination varied across treatments. For plants grown in control conditions, 21 VIs successfully discriminated genotypes carrying *HaHB11* from controls during the grain-filling stage, whereas, for the defoliation assay, only 14 VIs did the same (Supplementary Figure S5). For the waterlogging condition, the measurement carried out on the first date revealed differences in VI values, being E3 always significantly different from its control while E2 was only clearly discriminated from it in the control condition (Supplementary Figure S5). The VIs considering biomass, chlorophyll, and abiotic stress clearly differentiated transgenics from controls (Figure 7). Furthermore, PCA analyses for all treatments and all genotypes, showed the weights of the selected VIs, with vectors along PC1 and PC2 components far from 0, meaning that all of them are relevant to discriminate between genotypes. Besides, when a PCA was done per treatment, the Vis showed significant loadings, but they did not maintain the same clustering pattern (Supplementary Figure S5).

**Figure 7.**
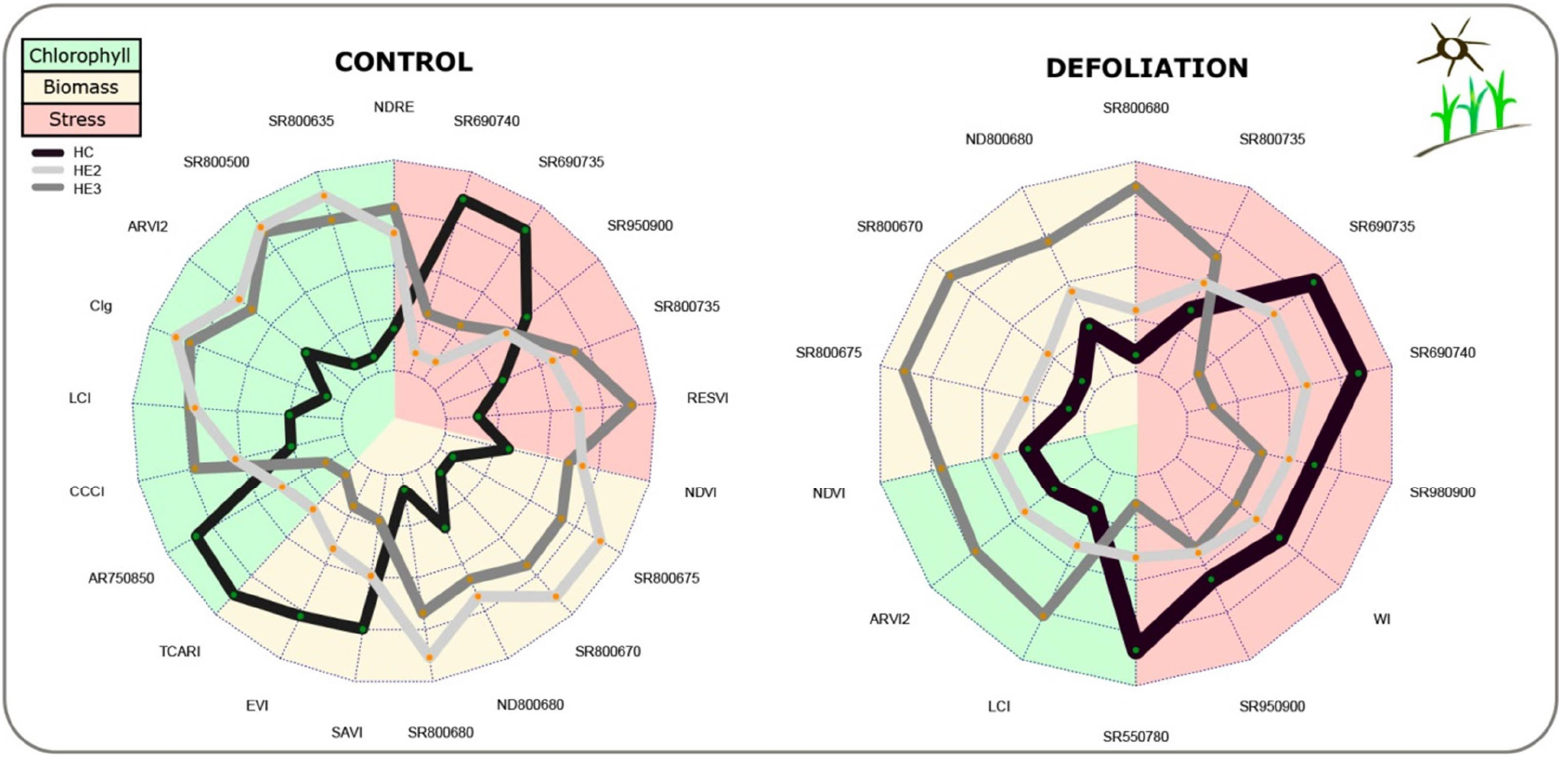
Vegetation indices computing spectral data are able to discriminate genotypes and treatments. Radar plots showing significantly different values of VIs between genotypes for standard and defoliation treatments. Black line: control genotype. Light grey and grey lines: E2 and E3 genotypes, respectively. Background colors indicate the applicability of the VIs. Green: chlorophyll, pink: water stress, light pink: biomass.

### The enhanced biomass, seed number, and grain yield exhibited by HaHB11 plants were robust and consistent traits across all the genetic backgrounds and tested conditions

Given the variety of growing conditions, environments, and genetic backgrounds in which the performance of the transgene *HaHB11* was tested, we performed a statistical analysis to determine the robustness of the differences in the evaluated traits as well as the relationship among them. Variance and multivariate analysis, including a principal component analysis (PCA), were carried out using all the data, considering inbreds, hybrids, greenhouse, and field assays in control and stress conditions (Supplementary Table S2 and Figure 8A).

**Figure 8.**
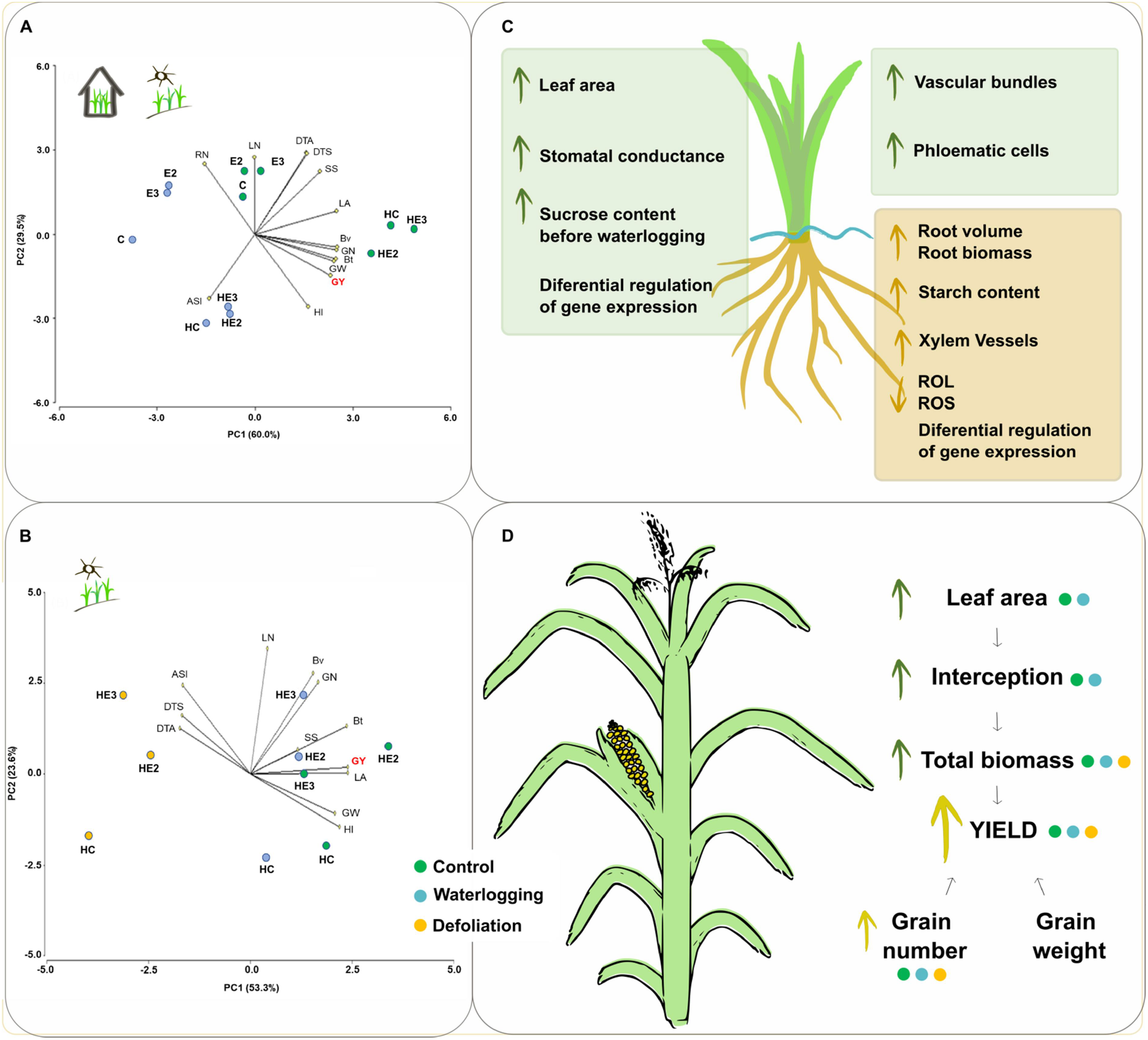
The beneficial effects conferred by *HaHB11* to maize plants are statistically robust. A and B. Principal component analyses (PCA) performed using all the data presented in this manuscript. C. Summary of the differential traits assessed between transgenic and control maize plants at the vegetative stage (V3) subjected to waterlogging. D. Summary of the differential traits assessed between transgenic and control maize plants at harvest after different treatments. Circles indicate (i) plants grown in control (blue) or waterlogging (green) conditions, and (ii) plants exposed to defoliation (orange).

Considering waterlogging effects, the PCA explained 89.5% of the variation produced by this condition early in development (Figure 8A). Within each water regime, hybrids had larger grain yield than inbreds and in the control condition, larger grain yield than under waterlogging stress. In general, both transgenic events, but particularly E3, had a larger grain yield than the control. Grain yield was associated with harvest index (acute angles between vectors) and did not respond to the variation in stem section, days to flowering, and ASI; vectors in right angle). Inbreds were located towards positive values of the PC2, with a higher number of roots and leaves and a longer vegetative period. Hybrids had enhanced harvest index (independently of the soil water condition) and longer ASI (particularly under waterlogging).

Aiming at knowing which traits are more related to the performance of HaHB11 plants in different conditions, an additional PCA was carried out considering only hybrids grown in the field subjected to stress caused by waterlogging or defoliation. The PCA explained 76.9% of the total variability. Hybrids carrying *HaHB11* always had a larger grain yield than the control under stressful conditions. Collectively, grain yield was tightly related to leaf area, total biomass, grain yield components, and harvest index; it had no relationship with leaf number and was negatively related to the extension of the vegetative period and the ASI. Within each growing condition, transformed genotypes tended to have larger grain numbers than the control, whereas the opposite trend was verified for individual grain weight (Figure 8B).

## Discussion

The expected second generation of transgenic plants is still absent in the market. The reasons for this are diverse, including the negative public perception of genetically modified crops and the fact that abiotic stress-tolerant crops do not represent a universal business since they are limited to target environments (Chan *et al*., 2020). However, the need for stress-tolerant crops remains actual and current due to global climate change and the increase of natural disasters. Although drought is still the major constraint for crop yield worldwide, flooding and severe storm episodes augmented their frequency with direct consequences for food and fuel production. Particularly, between 2006 and 2016, 65% of economic damage registered in crops was caused by abiotic stresses linked to excess water (FAO, 2017). To diminish the impact of such problems, breeders and biotechnologists work hard to obtain crops with improved behavior when exposed to abiotic and biotic stress factors. Usually, these efforts are not cooperative, and there is abundant scientific literature describing the research of stress-tolerant plants not tested in the field but only under controlled conditions or presenting very slight improvements (Sadras *et al*., 2020). Another important aspect of this research area is the slow but constant replacement of manual measurements by high-throughput phenotyping with modern, automated equipment that produces large databases and demands big data analysis. In this work, we presented the results of interdisciplinary research work, starting from molecular biology in the laboratory to spectral phenotyping in the field, performed to test the sunflower gene *HaHB11* as a potential tool to improve stress tolerance in maize.

The sunflower transcription factor HaHB11 has already been described as a transgene in the model Arabidopsis (Cabello *et al*., 2016) and maize plants (Raineri *et al*., 2019). In maize, transgenic plants were obtained in the ancient hybrid HiII (AxB) background and then backcrossed several times with the B73 line. When evaluated in greenhouse and field conditions under irrigation, these plants showed improved yield, mainly supported by a higher grain number than controls.

Maize is a species affected by flooding, mostly early in development (usually up to V2) and to a lesser extent in subsequent stages (Zaidi *et al*., 2004). Several works were dedicated to analyzing the effects of waterlogging, also called excess soil moisture (ESM) on contrasting genetic backgrounds (hybrids and inbreds). Such studies, applying varied waterlogging treatments, focused on physiological traits or/and molecular mechanisms. The more robust results indicated that the genotypes exhibiting early adventitious rooting, aerenchyma formation, a barrier to radial oxygen loss in roots, partial stomatal closure in leaves, and increase of NAD-ADH activity and starch accumulation in stem tissues were more tolerant to ESM than those that did not exhibit these traits (Zaidi *et al*., 2003). Notably, these attributes were common in induced hypoxia tolerance and allowed identifying associated QTLs (Abiko *et al*., 2012; Zaidi *et al*.., 2003). Increased ADH activity was a characteristic observed in adapted subtropical and tropical inbreds. Although the content of ethanol (the product of this enzyme) was higher in the susceptible genotypes, the ability to extrude it seemed to be increased in the tolerant ones (Zaidi *et al*., 2007). Tolerant and susceptible genotypes in advanced developmental stages differed in their ability to accumulate carbohydrates in stem tissues, the extension of the ASI, root porosity, and stomatal conductance (Zaidi *et al*., 2004). In agreement with these previous reports using different genetic backgrounds, *HaHB11* plants showed increased leaf area together with higher stomatal conductance, root length, and biomass than control plants after waterlogging treatments (Figures 1-4). Moreover, histological cuts of waterlogged roots evidenced tissue damage in the pith of control plants and an increased number of xylem vessels in the transgenics that could be associated with this stress response (Figures 1-5). Moreover, transgenic roots seemed to have a “tight” barrier to oxygen loss, compared to those of the wild type (Figure 5). After recovery, the transgenics exhibited an increase in nodal roots, stem surface, biomass, light interception, and chlorophyll, which resulted in an improved grain yield (Figures 1-5).

Regarding the mechanisms playing a role in waterlogging adaptation, inbreds showing susceptibility had a reduced dry matter translocation from source to sink tissues, which resulted in an inadequate grain filling (Kaur *et al*., 2021). Transgenic *HaHB11* plants, described here, accumulated more biomass and partitioned a larger part of it to grains, setting a higher grain number, resulting in increased yield compared to controls (Figure 5). An interesting question is if the waterlogging tolerance observed in several inbred genotypes was maintained in hybrids. It was reported that morpho-physiological traits differed between normal conditions and waterlogging and that hybrids were superior to parental lines under stress. Most of the characteristics associated with ESM tolerance in hybrids correlated positively with those of parental lines, but in normal moisture conditions, the effect of heterosis was more important than the contribution of the parental line (Zaidi *et al*., 2007).

In a more recent trial, different hybrids were tested in the V2 stage for their tolerance to waterlogging, evaluating similar parameters as in inbreds, after 7, 14, and 21 days of ESM. Although all the assessed hybrids showed a decrease in the evaluated traits, differences were detected between the tolerant and the susceptible ones (Kaur *et al*., 2021).

Regarding the molecular level, tolerant genotypes exhibited adaptive mechanisms enabling hypoxia tolerance. These plants had upregulated genes encoding enzymes participating in carbon metabolism and signal transduction, such as alcohol dehydrogenase, sucrose synthase, aspartate aminotransferase, NADP-dependent malic enzyme (Kaur *et al*., 2021). Notably, some of these genes were also differentially regulated in HaHB11 plants, albeit not always in the same sense (up or down, Figure 6). This contrasting result can be explained by the high turnover of genes observed after different waterlogging treatments and conditions. In transcriptome analyses performed with waterlogged maize plants, a huge variation in gene regulation was reported (Arora *et al*., 2017; Du *et al*., 2017; Rajhi *et al*., 2011). The only gene robustly regulated across all assays encoded a polygalacturonase (GRMZM2G037431, Li *et al*., 2019, Rajhi *et al*., 2011, Arora *et al*., 2017), and it was also upregulated in *HaHB11* transgenic plants.

Crop defoliation may recognize different origins, biotic like insect attack or abiotic as heavy rain, wind, or hail storms. In the trials described here, a summer storm produced severe defoliation (Figure 5). Although the serendipitous nature of the event, it allowed us to learn that *HaHB11* plants performed better than controls in response to such harmful conditions. The crop yield depends on the quality (i.e. size and activity) of the photosynthates source and the ability to transport assimilates to sink tissues. In maize, a consistent trend in seed dry weight was observed when assimilates during active grain filling were dramatically diminished by defoliation (Borrás *et al*., 2004). We hypothesized that because the storm occurred when the number of grains was almost already established, and transgenic plants set more grains than controls, the source-sink relationship may have been more affected among the former than among the latter. To corroborate this hypothesis, we developed controlled defoliation assays (Figure 5). As expected, the commercial hybrid, used as control, yielded more than all other hybrids in potential growing conditions; however, after a defoliation treatment, it was more penalized than the transgenic hybrids, indicating that a tolerant parental line in *HaHB11* plants contributed to a better performance in such condition. It is important to note that the influence of defoliation on crop yield depends on its timing and severity. During vegetative stages, it may have no or little effect, whereas even a mild hail can produce a reduction on grain yield of 30% or more when it occurs from the start of the critical period onwards (Battaglia *et al*., 2018), depending upon de relative impact on light interception efficiency (Borrás *et al*., 2004; Cerrudo *et al*., 2013).

Described differences among genotypes across treatments and environments were confirmed by multivariate analysis and assessed through vegetation indices (VIs) indicative of variations in canopy spectral reflectance. Rather than selecting the best VI for describing each evaluated trait (García-Martínez *et al*., 2020), we opted for a joint analysis of 29 VIs to capture differences among genotypes in a single environment (Arias *et al*., 2021). Although genotypes could be differentiated, it was not the same set of VIs that allowed their discrimination across treatments. On the one hand, this response is indicative of the capacity of VIs to track the fast changes in plant metabolism in response to the environment (Reynolds *et al*., 2021). On the other hand, described shifts in the way VIs ranked genotypes along the cycle alert on the need for further research aimed to understand the interrelation between spectral data and differential gene expression among genotypes.

## Conclusions

In current research, we demonstrated the advantage of maize genotypes transformed with *HaHB11* to withstand transient episodes of ESM that take place early in the cycle. This response is probably linked to their improved root system because the negative effects of ESM in mentioned stages usually affect roots more than shoots (de San Celedonio *et al*., 2017), and we observed that starch content accumulated in leaves and decreased in roots of the control genotypes whereas the opposite trend occurred in genotypes transformed with *HaHB11*. The latter is indicative of an active metabolism despite the stressful anaerobic condition. Being the recovery of roots lower than that of shoots, plants bearing *HaHB11* may be in better conditions than the controls to withstand subsequent stressful scenarios along the cycle (e.g. defoliation, enhanced evaporative demand), which are not uncommon among field-grown plants.

## Materials and Methods

### Plant material and growth conditions

Greenhouse assays: maize plants of different genotypes (B73 lines and B73 x Mo17 hybrids) were grown during 2017 and 2021 at the Institute of Agrobiotechnology, located at Santa Fe (31°38’17.1”S, 60°40’01.8”W). Plants were cultivated in 45 L pots under long-day photoperiod (16/8 h light/dark cycles), with daily temperatures fluctuating between a mean minimum of 10°C and a mean maximum of 40°C.

Field trials: experiments were performed during 2017 and 2020 at the IAL, on a sandy soil of 2 m depth with low water-holding capacity and an organic ‘A’ horizon of 15 cm. Evaluated germplasm included the following genotypes: controls (null segregants) and transgenic lines (E2 and E3) of B73 lines (transformed HiII backcrossed 3-4 times to B73, experiments 1 and 2) as well as B73 × Mo17 hybrids (experiment 3) and commercial F1 hybrid DK 72-10®(all experiments). The sowing date took place at the beginning of November using a single stand density of 9 plants m^-2^. Genotypes were distributed in a completely randomized design with three replicates. Each plot had three rows of 1.25 m and 0.5 m between rows. Plots were drip-irrigated along the whole cycle to keep the uppermost 1 m layer at field capacity and were fertilized with N (180 kg ha^-1^ at sowing and 180 kg ha^-1^ at tasseling) and P (100 kg ha^-1^ at sowing). Plots were kept free of weeds, insects, and diseases. Daily mean temperature (in °C) and incident solar radiation (in MJ m^-2^ day^-1^) were obtained from a nearby meteorological station. All experiments were carried out after obtaining the corresponding authorization from the CONABIA (National Committee of Biotechnology) and INASE (Seeds National Institute).

### Waterlogging treatments

In greenhouse experiments, plants (all the tested genotypes) with three expanded leaves (V3) were placed in a plastic pool. At the beginning of the day, enough water was added to cover half of the pot height. At midday, the water level raised until 1 cm above the ground for two weeks.

In field assays, the pots were placed in pools. At the V3 stage, plants were waterlogged during 14 days, keeping the water level 1 cm above the ground. After the treatment plants from all genotypes were placed at the same time in the experimental field and grown under irrigation until the end of the life cycle.

### Defoliation assays

In the field trial, the first defoliation episode occurred 11 days after silking. The second and third experiments were carried out in the field and greenhouse, respectively, by manually defoliating plants, eleven days after silking leaving only the ear leaf and the leaf immediately above it. All ribs were kept, emulating the field defoliation.

### Plant and crop phenotyping

Measurements were performed on five plants per genotype (greenhouse) or six plants from the central row of each plot (field) which were tagged at V3, as previously described (Raineri *et al*., 2019). Dates of ASI were registered for all plants. At silking, the total number of fully expanded leaves and total plant leaf area were measured. Individual leaf area was computed as in Equation 1 (Montgomery, 1911)

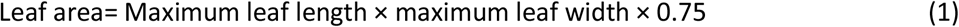

All tagged plants were oven-dried for estimation of total aerial plant biomass. Grain yield was expressed on a 14.5 % wet basis. Harvest index (HI) was estimated as the quotient between grain yield and plant biomass (on a dry basis).

### Stomatal conductance

Stomatal conductance was measured with a porometer (Decagon® SC-1). All measures were taken at midday.

### Carbohydrate and chlorophyll contents

Starch, sucrose, glucose, and protein contents from roots and leaves of at least 4 plants were assessed as previously described (Cabello *et al*., 2016). Chlorophyll content was determined either by acetone extraction (Raineri *et al*., 2019) or by using a specific chlorophyll meter device (Cavadevices®, https://cavadevices.com/archivos/FOLLETOS/Clorofilio.pdf).

### Allometric measurements during waterlogging stress in roots

At least four plants were harvested and the roots were washed. Total adventitious roots, root length and root were quantified. Root volume was assessed by a volumetric method: 80 ml of deionized water (V_W_) was placed in a tube; then, the root system was completely submerged and the volume of water plus the root (V _W+R_) was measured. The root volume (V_R_) was calculated as follows: V_R_ = V _W+R_ -V_W_

### Radial oxygen loss

The radial oxygen loss in roots was measured as described (Watanabe *et al*., 2017). Four plants per genotype were selected after the waterlogging treatment. All the adventitious roots were removed, except one of 10-14 cm length. Plants having a single nodal root were placed in a pot with methylene-blue solution and photographed after 1 hour of incubation.

### NBT staining

Roots were collected and placed in a solution containing NBT 0,1 mg/ml in 25 mM Hepes pH 7,6 and 0,05% Triton X-100. The samples were vacuum-infiltrated for 15 minutes and incubated for an additional hour at 37°C.

### RNA isolation and expression analyses by real-time RT-PCR

Total RNA for real-time RT-PCR was isolated from maize leaves or stems using Trizol® reagent (Invitrogen, Carlsbad, CA, USA) and real-time qPCR was performed using an Mx3000P Multiplex qPCR system (Stratagene, La Jolla, CA, USA) as described before (Raineri *et al*., 2019). Primers used are listed in Supplementary Table S1.

### Histology

Histology of the cross-sections was carried out as previously described (Cabello *et al*., 2016) and stained with safranine fast-green. The xylem and pith area were assessed using the free software ImageJ (Schneider *et al*., 2012). For lignin content, the cross-sections were evaluated using an epifluorescence microscope.

### Remote sensing analyses

Canopy spectral reflectance was measured using a compact shortwave NIR spectrometer (Ocean Insight). The instrument is sensitive to 1024 wavelengths in the range from 632 nm to 1125 nm with an optical resolution of 3 nm at full-width half-maximum. In situ measurements were performed between 10:00 and 14:00 h ART time (UTC 03:00), with the instrument positioned at a nadir view 50 cm above the surface. The upwelling light reflected from a 50 cm x 50 cm white reference material with 99% reflectance, was recorded before each canopy measurement allowing data acquisition during variable sky conditions. The integration time was adjusted to avoid saturation of the white signal and each measurement was the average of five successive scans. The measurements were homogeneously distributed over the plot to reduce border effects. Measurements were collected on 01/17/2020 (vegetative stage), 01/28/2020 (flowering); and 02/18/2020 (grain-filling stage). A typical outlier control based on standard deviation was implemented on each canopy spectral reflectance raw data.

Twenty-nine vegetation indices were selected based on the range of available wavelengths and their applications. The selected indices, their formulas, and type of applications are shown in Supplementary Figure S1.

Each vegetation index was evaluated for each treatment per genotype combination. Only those that differed significantly (P<0.05) between genotypes are discussed. Data analysis was conducted in R using the aov function and the post-hoc test was performed using the agricolae-package (Mendiburu, 2010).

### Statistical analyses

A t-test was used for the comparison of genotypes evaluated in the greenhouse experiment, whereas ANOVA was used to assess the effect of treatments (control, waterlogging, and defoliation), genotypes (line or hybrid, control or transgenic), and their interaction on the evaluated traits in greenhouse and field experiments. Differences across means were analyzed by a Tukey test (Supplementary Table S2). Principal components analyses (PCA) were used to evaluate the correlation among traits for the different genotypes and experiments, as well as for the vegetation indices (Figure 8 and Supplementary Figure S1).

### Accession numbers

Accession numbers of the genes evaluated in this work are available in Supplementary Table S1

## Acknowledgements

We thank Silvia Lede for her professional assistance in the procurement of CONABIA and INASE (Ministry of Agriculture) permits. We are very grateful for the technical assistance provided by Mr. Manuel Franco.

## Notes

**Funding information** This work was supported by Agencia Nacional de Promoción Científica y Tecnológica (PICT 2015-2176), and CONICET. LC is an undergraduate student at Universidad Nacional del Litoral. JR and NR are Fellows of CONICET, MEO and RLC are career members of the same institution. MP is Professor at the National University of Rosario.

